# Post-translational modifications on protein VII are important during the early stages of adenovirus infection

**DOI:** 10.1101/2024.08.21.609089

**Authors:** Edward A Arnold, Julian R Smith, Katie Leung, Daniel H Nguyen, Laurel E Kelnhofer-Millevolte, Monica S Guo, Jason G Smith, Daphne C Avgousti

## Abstract

Due to the importance of post-translational modification (PTM) in cellular function, viruses have evolved to both take advantage of and be susceptible to such modification. Adenovirus encodes a multifunctional protein called protein VII, which is packaged with the viral genome in the core of virions and disrupts host chromatin during infection. Protein VII has several PTMs whose addition contributes to the subnuclear localization of protein VII. Here, we used mutant viruses that abrogate or mimic these PTMs on protein VII to interrogate their impact on protein VII function during adenovirus infection. We discovered that acetylation of the lysine in positions 2 or 3 (K2 or K3) is deleterious during early infection as mutation to alanine led to greater intake of protein VII to the nucleus and enhanced early gene expression. Furthermore, we determined that protein VII is acetylated at alternative residues late during infection which may compensate for the mutated sites. Lastly, due to the role of the early viral protein E1A in viral gene activation, we investigated the interaction between protein VII and E1A and demonstrated that protein VII interacts with E1A through a chromatin-mediated interaction. Together, these results emphasize that the complexity of virus-host interactions is intimately tied to post-translational modification.

**Importance:** Adenoviruses are a ubiquitous human pathogen that cause a variety of diseases, such as respiratory infections, gastroenteritis, and conjunctivitis. While often viewed as a self-limiting infection in healthy individuals, adenoviruses are particularly harmful for immunocompromised patients. Here, we investigate the functional role of post-translational modifications (PTMs) on an essential adenovirus core protein, protein VII, describing how they regulate its function during early and late stages of infection. Our study focuses on how specific PTMs on protein VII influence transcription, localization, and interactions with other proteins, highlighting how PTMs are employed by viruses to alter protein function.

## Introduction

Chromatin is a complex of DNA and histone proteins that serves to compact the host genome and regulate gene expression (1). Histones are heavily marked with post-translational modifications (PTMs) such as phosphorylation, acetylation, and methylation, among others, that impact DNA accessibility through compaction. For example, acetylation of histone H3 at lysine 18 (H3K18ac) is associated with open chromatin and active gene expression (2, 3). In contrast, methylation at lysine 9 (H3K9me) is commonly associated with repressed chromatin (2, 3).

Phosphorylation of histones is frequently involved in the activation of many signaling pathways such as phosphorylation of H3S10 (H3S10ph) during mitosis (2, 4–6). Similar to histones, protamines are small arginine rich basic proteins that, in mammals, serve to compact the paternal genome within sperm. The paternal genome is compacted within the restricted space of sperm cells supporting the notion that protamines are more efficient at compacting the genome than histone proteins (7, 8). Upon fertilization, protamines are extensively modified, which is hypothesized to reduce their high charge and promote their replacement with histones (7). Thus, PTMs are an essential aspect of the relationship between DNA and histones or histone-like-proteins.

Adenovirus is a double-stranded DNA virus that encodes a core protein, called protein VII, which is bound to the viral genome within the virion and has been described as a histone or protamine-like protein (9–11). Protein VII is a small, basic protein of 198 amino acids that is initially expressed as a precursor, preVII, before a viral protease cleaves the first 24 amino acids resulting in the packaged mature protein VII (12–14). Protein VII is essential in that although protein VII-null virions can be made, the incoming virions remain trapped in the endosome during viral entry and are unable to establish an infection (15). In a wild-type infection, protein VII is delivered with the genome to the nucleus (16, 17). Protein VII foci, visualized by immunostaining, are commonly used as a marker for incoming viral genomes. These incoming genomes can be visualized as foci that persist up to 10-12 h post-infection (p.i.) (17–20).

Additionally, chromatin immunoprecipitation of protein VII showed that protein VII associates with the viral genome up to 10-12 h p.i. and is slowly removed over time, independent of transcription or DNA replication (17, 21–25). In contrast, other reports showed that protein VII foci decreased with the onset of transcription of early viral genes and that protein VII foci persisted when transcription was inhibited or in the absence of early viral gene E1A (19). E1A binds to the host genome and facilitates transitioning the cell into a viral ‘S phase’, and importantly activates transcription of other early viral genes (26), among other functions. Protein VII also protects the incoming viral genome from recognition by the host DNA damage response (27, 28), indicating that the function of protein VII on the incoming viral genomes is more complex than gene expression regulation. Nevertheless, how the presence of VII impacts the onset of viral gene expression is unclear.

Protein VII was initially hypothesized to act as a transcriptional repressor because remodeling of the incoming viral genome to deposit histones and initiate transcription temporally coincided with the loss of protein VII (19, 25). A recent study used chromatin profiling techniques to map protein VII on the early viral genome and found that protein VII forms a repeating complex with DNA, termed adenosomes, that are reminiscent of host nucleosomes (29). Schwartz *et al* found that lower amounts of protein VII on a viral gene, for example early genes, was correlated with transcription, supporting the idea of protein VII as a repressor (29). In contrast, other studies proposed that protein VII activates transcription (30–32), likely through its interaction with chromatin factor TAF-Iβ/SET that promotes remodeling of the incoming viral genome for gene expression (18, 22). Further, it was hypothesized that protein VII directly binds and recruits newly synthesized E1A to the viral genome to drive early gene transcription (25), though the order of events has yet to be determined.

Interestingly, protein VII also binds fully formed nucleosomes (33), which suggests that the interaction of protein VII with DNA or newly deposited nucleosomes on the viral genome may be more nuanced than previously thought. During late stages of infection, protein VII is expressed with other late genes to high levels. At this time, protein VII localizes both to viral replication compartments (VRCs) for assembly into nascent virions and to host chromatin, where it is thought to bind nucleosomes directly with the aid of host factors SET and HMGB1 (33–35). The presence of protein VII on the host genome leads to disruption of the cell cycle (34), though impact of protein VII on host transcription has not yet been elucidated.

Much like histones and protamines, protein VII is also post-translationally modified. The mature protein contains two acetylation sites (K2 or K3 and K24) and three phosphorylation sites (T38, T48 or T50, and S159) (33). Additional phosphorylation (S19) and acetylation (K20) sites were also identified on the N-terminal precursor fragment (33). Wild-type mature protein VII localizes to host chromatin upon ectopic expression; however, if all identified PTM sites are mutated to alanine to abrogate modification, protein VII no longer localizes to chromatin but to the nucleolus (33). Further analysis suggested that acetylation of either K2 or K3, which appear to be redundant, is critical for host chromatin localization. As such, mutation of both lysine residues to alanine in ectopically expressed protein VII phenocopied the mutation all PTM sites and resulted in nucleolar localization, whereas mutation of either lysine residue to the acetyl mimic glutamine recapitulated the wild-type localization to chromatin (33). These data support a model for the distribution of protein VII during infection wherein unmodified protein VII localizes to viral genomes in VRCs to be packaged while modified protein VII localizes to host chromatin. Consistent with this model, mass spectrometry on purified virions identified no acetylation on protein VII or phosphorylation at T72 or S79 (33, 36, 37), suggesting that PTMs may contribute to protein VII’s ability to distinguish between viral and host chromatin.

Here, to investigate the function of PTMs on protein VII during infection, we generated mutant viruses with point mutations in protein VII. We demonstrate that preventing acetylation of K2 or K3 by mutating both residues to alanine led to enhanced nuclear entry of the viral genome which causes earlier onset of expression of the early viral gene E1A. Interestingly, this acceleration of early gene expression caused by mutating protein VII had no impact at later stages of infection. Further, we found that mutated protein VII is still acetylated during infection, likely at alternate residues. To elucidate the dynamics of early infection, we discovered that protein VII and E1A interact in a chromatin-dependent manner, adding a new dimension to our understanding incoming viral genome remodeling and the onset of early transcription. Together, our findings establish that modifications on protein VII are critical for early viral entry and gene expression.

## Results

### Post-translational modifications on protein VII contribute to nuclear entry

Due to protein VII’s role in establishing infection, we hypothesized that PTMs on protein VII may impact early stages of infection. To test this, using recombineering (35, 38) we created replication-competent E3-deleted human adenovirus type 5 (HAdV-5) mutants with point mutations and a C-terminal HA tag on protein VII. We generated three mutant viruses in which (1) all five PTMs sites on protein VII were mutated to alanine (ΔPTM), (2) only the second and third lysine residue were mutated to alanine (K2AK3A), or (3) the third lysine residue was mutated to the acetyl mimic glutamine (K3Q) (Fig 1). We then infected A549 cells with these viruses and visualized incoming genomes by immunostaining for HA at early time points of infection. To ensure comparable infection dynamics, we synchronized infection by cold incubation prior to initial infection (see Methods). We also immunostained for the early viral protein E1A to monitor the onset of early viral gene expression (Fig 2A). Next, we quantified the integrated density of HA staining within the nucleus of infected cells (Fig 2B). At 2 h p.i., the majority of HA-positive foci localized to the cytoplasm in all samples, with very little positive staining in the nucleus (Fig 2A and 2B). This is somewhat unexpected since adenovirus genomes have been reported to reach the nucleus by approximately 45 minutes after infection (39, 40); however, since our system uses an HA-tagged protein VII in an E3-deleted virus with cold synchronization, these differences could account for a delay in kinetics. At 4 h p.i., we observed HA-positive foci both in the nucleus and the cytoplasm, and the nuclear localization of ΔPTM and K2AK3A foci were significantly greater than WT (Fig 2A and 2B). At 6 h p.i., this difference became more pronounced, with even more foci in the nucleus upon infection with the K2AK3A and ΔPTM compared to the other viruses. By 8 h p.i., the pattern remained the same for all four viruses, with the K2AK3A virus infected cells containing the most nuclear foci; however, the HA signal became more diffuse at this time, suggesting that protein VII may no longer be associated with viral genomes. By quantification, we observed that the WT and K3Q viruses had similar patterns of nuclear entry, meaning that the distribution of foci at 4 h p.i. were very similar, suggesting that the K3Q virus is phenotypically similar to the WT. Interestingly, the K2AK3A and ΔPTM viruses also had comparable patterns at 4 h p.i. with a significantly higher proportion in the nucleus compared to the other two viruses, suggesting that the mutated lysine residues on protein VII, or their acetylation, are important for nuclear entry of the viral genome.

**Figure 1:**
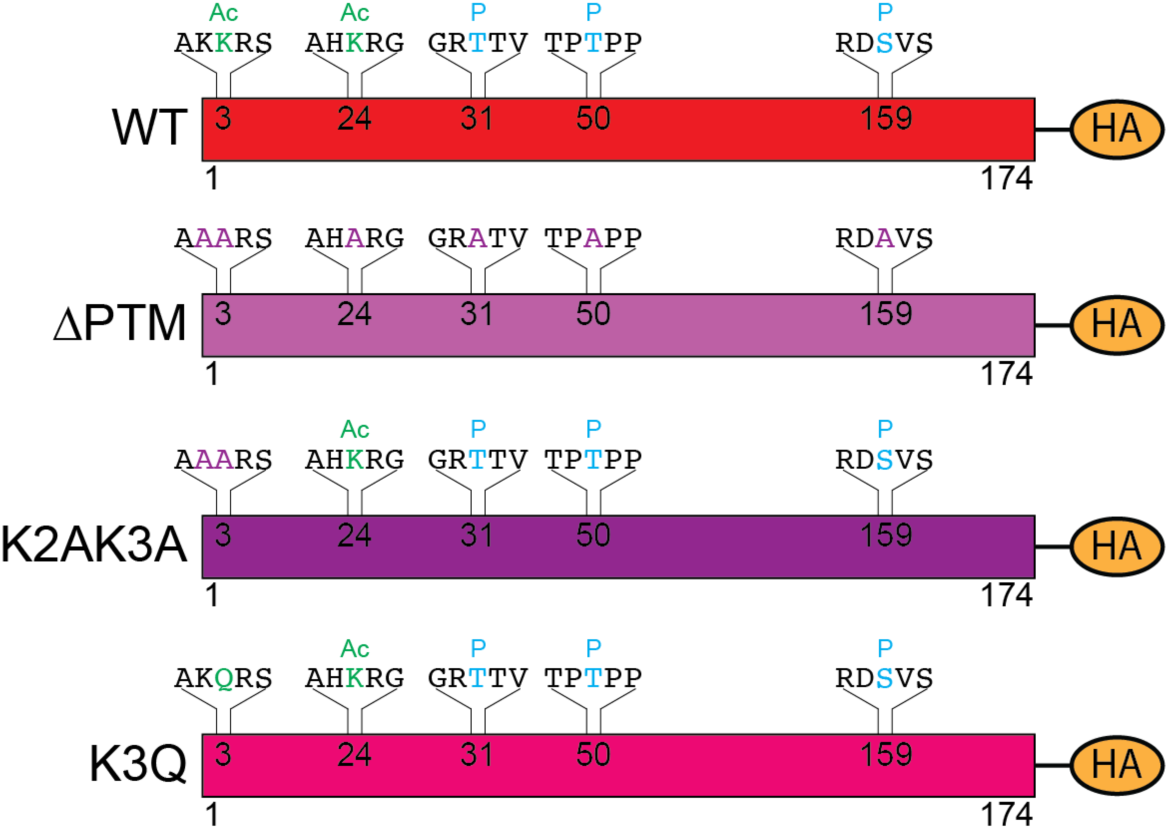
Schematic of mutant virus design. Schematic of protein VII from wild-type (WT) and mutant viruses. WT mature protein VII (top) with a C-terminal HA tag is shown. The amino acid sequence and corresponding residue positions are indicated. Amino acid changes and their predicted effects on modifications are shown for mutant viruses below. Ac is acetylation in green, P is phosphorylation in cyan, and alanine substitutions are in purple.

**Figure 2:**
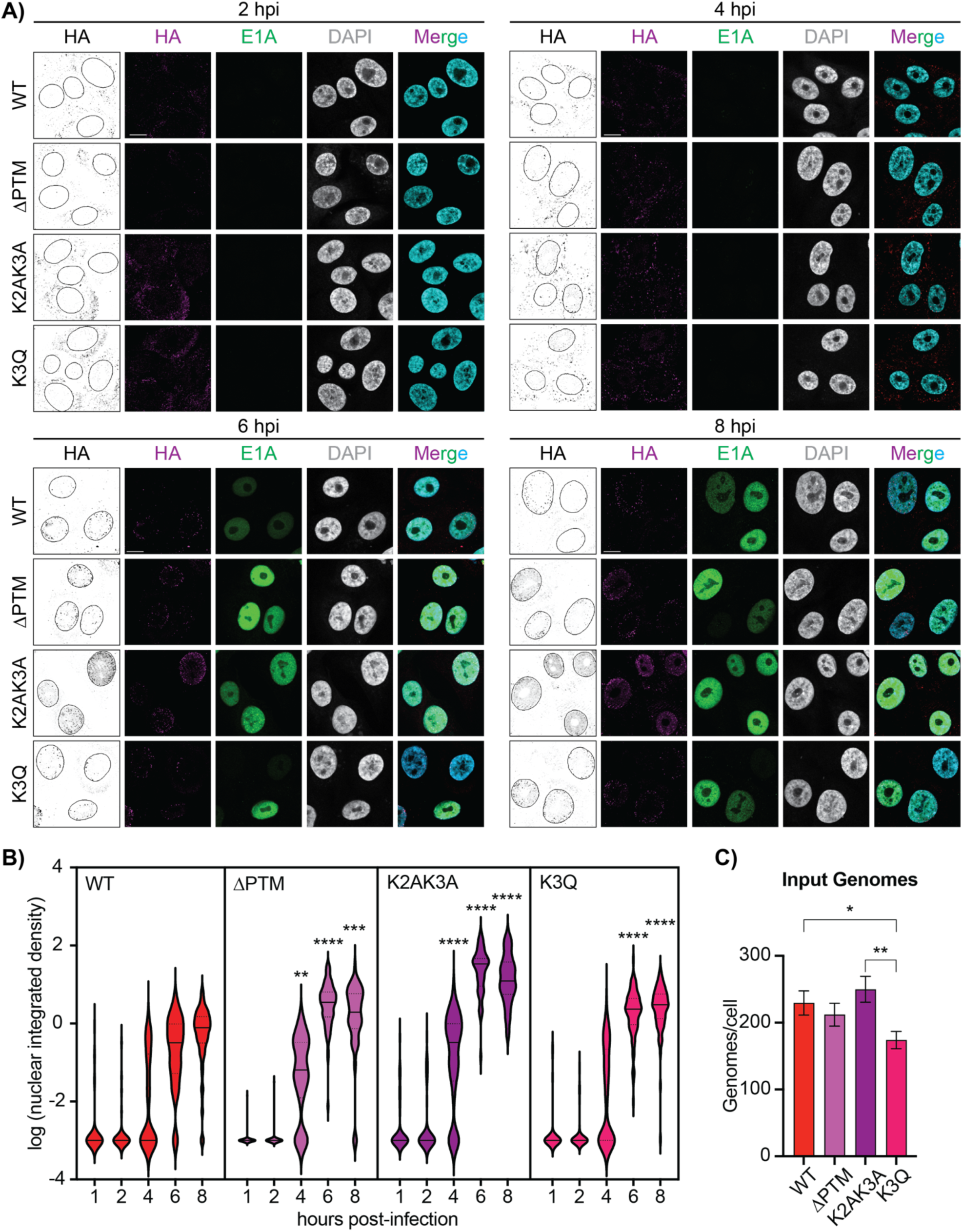
Post translational modifications on protein VII are important for viral entry into the nucleus. A) Representative immunofluorescence images of A549 cells infected with WT and mutant viruses at 2 (top left), 4 (top right), 6 (bottom left), and 8 (bottom right) h p.i. as indicated, showing HA in magenta, E1A in green, and DAPI in grey (cyan in merge). Additional panels of HA staining in black and white and nuclei outlined are provided for better visualization. Scale bar is 10 µm. B) Quantification of the integrated density of HA staining within the nucleus for each time point. N>30 nuclei for each virus at each time point. ** is p<0.01, *** is p<0.001 **** is p<0.0001 by one-way ANOVA with Dunnett’s multiple comparisons test for each condition compared to WT. C) Quantification of input viral genomes per cell by qPCR of infected A549 cells with indicated viruses at MOI of 25 pfu per cell. Input genomes are presented as mean and error bars represent standard deviation. * is p<.05, ** is p<.01 by two-way ANOVA with Tukey’s multiple comparisons test. n=3 biological replicates.

We also observed that the K2AK3A virus had a higher integrated density plateau in our quantification analysis. We hypothesized that this may be due to differences in the particle to pfu ratio of our stock viruses. To ensure comparable amounts of virus were used in each condition, we performed qPCR to determine the average number of genomes per cell and found that the K2AK3A had the highest (Fig 2C) and that the WT and K2AK3A viruses were both significantly higher than the K3Q virus (Fig 2C), suggesting more viral genomes were present. Nevertheless, this slightly higher plateau does not account for the significantly greater foci detected in the nuclei of cells infected with the K2AK3A and ΔPTM viruses. Furthermore, we observed that E1A became detectable at 6 h p.i. and increased in intensity by 8 h p.i. We found both a greater number of E1A positive cells as well as more intense E1A signal in the K2AK3A and ΔPTM virus infected cells compared to the WT and K3Q virus infected cells at 6 h p.i. These results suggest that the increased nuclear entry observed by immunofluorescence also leads to increased expression of E1A. Taken together, these results suggest that mutation of the K2 or K3 residues of protein VII enhances viral genome entry into the nucleus.

### K2AK3A mutations in protein VII enhance early gene expression

Because of the increase in E1A positive nuclei in the K2AK3A and ΔPTM infected cells, we hypothesized that transcription may also be starting earlier with mutation of K2 and K3 on protein VII. To investigate this, we performed infections over a time course with our mutant viruses and measured viral mRNA by RT-qPCR at 0.5, 1, 1.5, 2 and 4 h p.i. and protein expression by western blotting at 4, 6 and 8 h p.i. (Fig 3). We found an increase in relative abundance of *E1A* mRNA in the K2AK3A infection, and to a lesser extent in the ΔPTM infection, at all early time points tested (Fig 3A). These differences were noticeable by 1 h p.i. and the E1A mRNA remained consistently higher for the K2AK3A and ΔPTM virus until 4 h p.i. when the difference reached statistical significance for both the K2AK3A and ΔPTM viruses compared to WT. The K3Q virus produced mRNA levels similar to those of the WT virus throughout the time course, consistent with the idea that K3Q is a mimic of acetylation which is present on WT protein VII, while the K2AK3A and ΔPTM viruses exhibited earlier expression, consistent with earlier entry into the nucleus.

**Figure 3:**
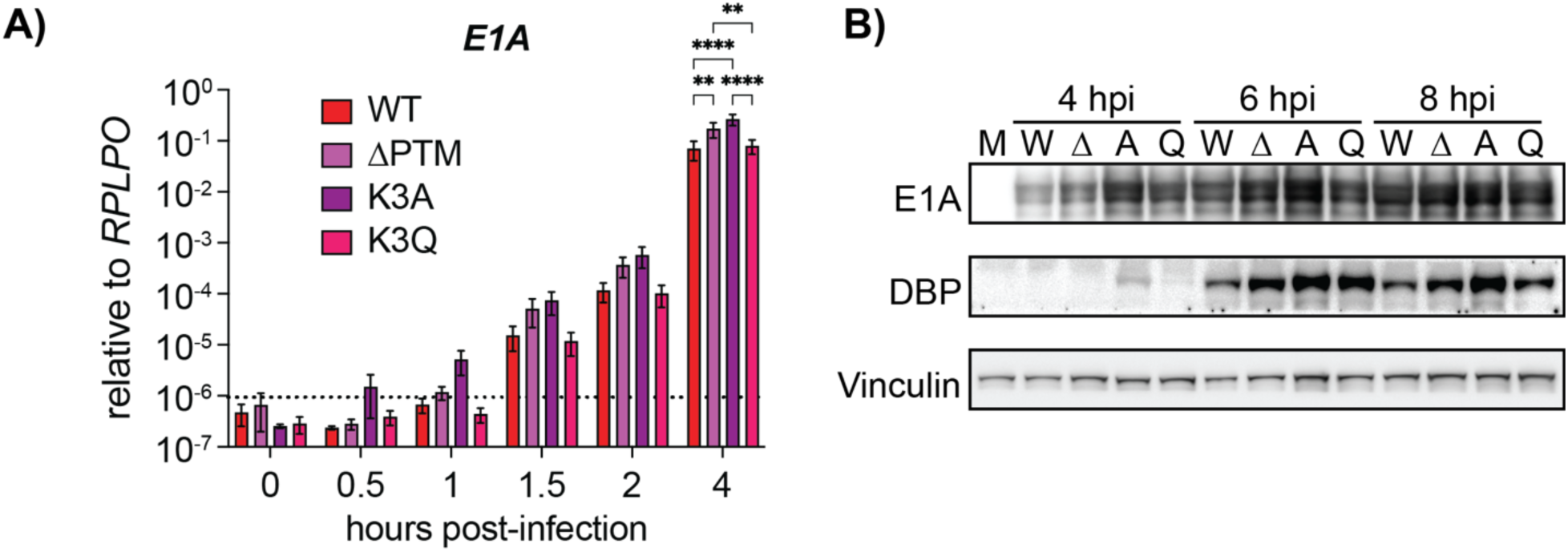
Mutation of lysine 2 and 3 on protein VII enhances early gene expression. A) E1A mRNA levels measured by RT-qPCR of A549 cells infected with corresponding viruses, cold synchronized for 1 h on ice, and then collected at the indicated time points. mRNA levels are presented as mean, error bars represent standard deviation. **, p<0.01, and ****, p<0.0001 by two-way ANOVA with Tukey’s multiple comparisons test, n = 3 biological replicates. Dotted line represents the limit of reliable detection by qPCR. B) Representative western blots of A549 cells infected with the indicated viruses and harvested at 4, 6, and 8 h p.i. Blots were stained for viral early proteins E1A and DBP and vinculin as a loading control. M is mock, W is WT, Δ is ΔPTM, A is K2AK3A, and Q is K2Q. n = 3 biological replicates.

We next examined the expression of early viral proteins by western blot and found that E1A protein levels in the K2AK3A virus infection were greater compared to the other viruses at 4 and 6 h p.i. (Fig 3B). By 8 h p.i., E1A expression appeared to reach similar levels across all four viruses. We also probed for another early viral protein, E2A, more commonly referred to as DNA binding protein (DBP), which mirrored the E1A results. At 4 h p.i., DBP was only visible in the K2AK3A virus, and was more intense at 6 and 8 h p.i. compared to the other viruses. Taken together, these findings suggest that mutation of these lysine residues on protein VII to alanine results in greater numbers of genomes entering the nucleus, leading to increased early gene expression.

### Post translational modifications on protein VII do not impact later stages of infection

Due to the effects of protein VII mutations on early viral gene expression, we next investigated their impact at later stages of infection. To measure this, we examined genome replication, protein production, and infectious progeny production. We performed qPCR to measure relative genome amounts and observed no statistically significant difference in genome replication across the four viruses (Fig 4A). Similarly, there were no observable differences in viral protein levels between the four viruses (Fig 4B). Lastly, we measured the amount of viral progeny produced during infection by plaque assay. All four viruses had similar titers at the three time points tested, suggesting that these mutations did not impact progeny production (Fig 4C). Despite the enhanced early gene expression upon K2AK3A mutation, and to a lesser extent in the ΔPTM virus, our data show that all four viruses appear to function similarly by later stages of infection.

**Figure 4:**
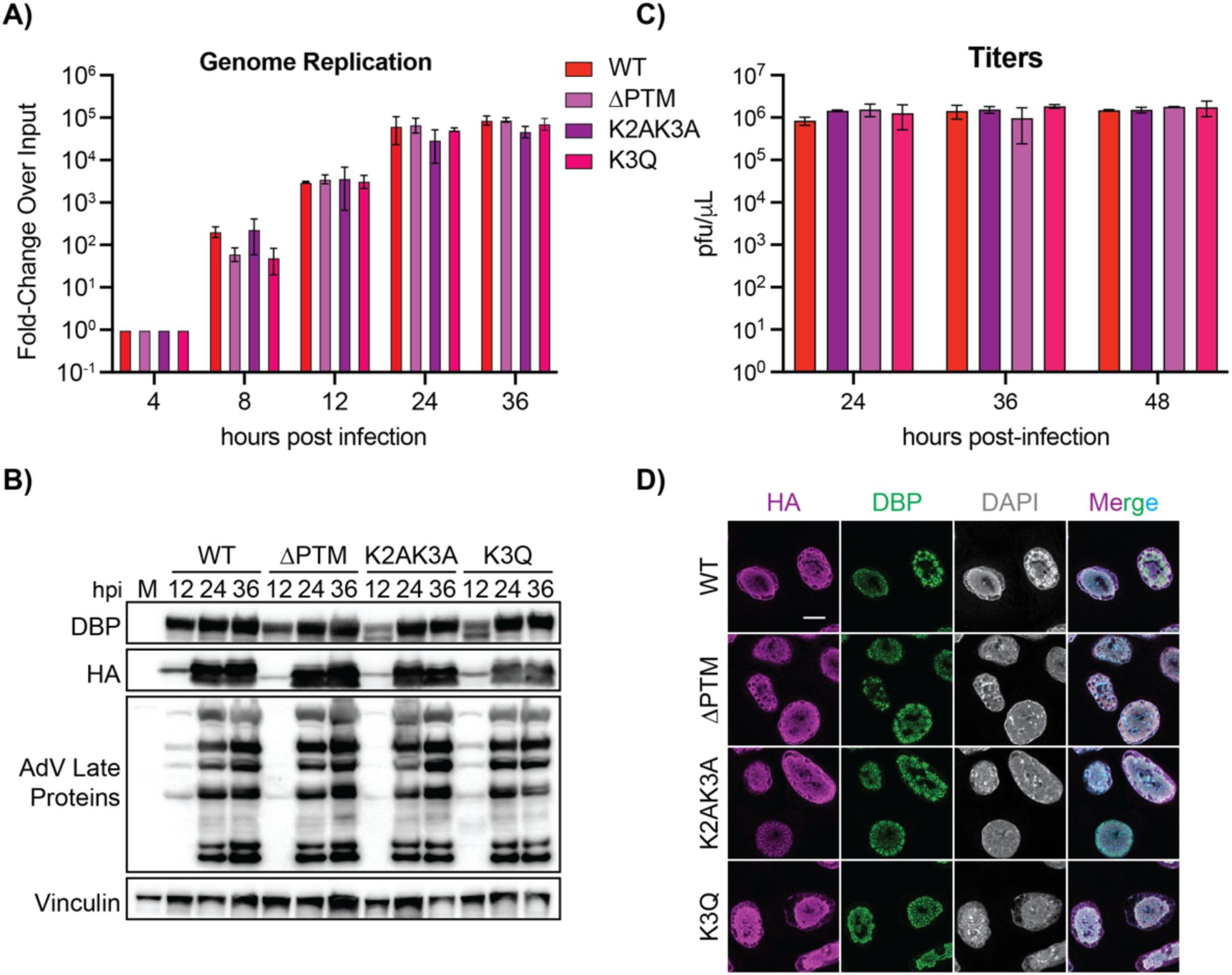
Mutation of modified residues on protein VII does not significantly impact genome replication, late protein production, or infectious progeny production. A) Genome quantification as measured by qPCR of A549 cells infected with indicated viruses at 4, 8 12, 24, and 36 h p.i., depicted as mean with error bars representing standard deviation. n = 3 biological replicates. No significance by two-way ANOVA with Tukey’s multiple comparisons test. B) Representative western blot of A549 cells infected with indicated viruses at time points as indicated. Blots were stained for protein VII (HA), early viral protein DBP, late viral proteins, and vinculin as a loading control. n = 3 biological replicates. No significance by two-way ANOVA with multiple comparisons. C) Infectious progeny quantification of the indicated viruses upon infection of A549 cells at 24, 36, and 48 h p.i. Titers are presented as mean with error bars representing standard deviation. n = 3 biological replicates. No significance by two-way ANOVA with Tukey’s multiple comparisons test. D) Immunofluorescence images of A549 cells infected with WT and mutant viruses at 14 h p.i. HA is magenta, DBP is green, DAPI is gray (cyan in merge). Scale bar is 10µm. n>30 nuclei for each virus.

PTMs on protein VII impact localization of the ectopically expressed protein such that WT and K3Q protein VII localized to chromatin, while ΔPTM and K2AK3A protein VII localized to the nucleolus (33). To assess the impact of these PTMs on localization during infection, we infected A549 cells with the WT and mutant viruses and examined cells at 14 h p.i. by confocal microscopy. We immunostained for HA to visualize protein VII localization, DBP to visualize infection progression, together with DAPI. We found that WT protein VII had a dispersed nuclear localization, forming large puncta throughout the nucleus consistent with previous reports (Fig 4D). Surprisingly, there was no clear difference in protein VII localization across the mutant viruses. In fact, contrary to ectopic expression (33), the ΔPTM and K2AK3A mutants did not localize to the nucleolus and were equivalent to WT and K3Q. Thus, we conclude that the localization of newly synthesized protein VII during the later stages of infection is not impacted by mutation of protein VII at these specific residues.

### The ΔPTM mutant is alternatively acetylated in infected cells

Next, because our mutant viruses had no major phenotypic changes in comparison to the WT virus at later stages of infection, we hypothesized that ΔPTM and K2AK3A may be modified at different nearby residues to compensate for the mutations. While the structure of protein VII has not been solved, we used the protein prediction software I-TASSER to model protein VII (41, 42). This analysis produced a structure of a bundle of seven α-helices (red, Fig 5A), with an eighth helix for the N-terminal precursor fragment (magenta, Fig 5A). The remaining lysine residues are K20 in the precursor fragment and K73 and K75 of the mature protein (K97 and K99 of preVII) (dark gray, Fig 5A). In the predicted structure, K20 within the precursor fragment is in close physical proximity to the known acetylated lysine residues K2 and K3 (K24 and K25 of preVII, green Fig 5A). K73 and K75 (dark gray, Fig 5A) are also predicted to be on the end of a helix, making them accessible for acetylation. Due to the locations of lysine residues within the predicted structure of protein VII, we hypothesized that in the absence of the preferentially modified lysine residues (K2, K3, and K24), an acetyl transferase enzyme may modify other nearby lysine residues. To test this, we performed an immunoprecipitation (IP) of protein VII from WT and ΔPTM infected cells at 24 h p.i. when protein VII is expressed to high quantities and probed with a pan lysine-acetyl antibody. We found that both the WT and ΔPTM viruses showed the double banding pattern typical of the pre- and mature protein VII (Fig 5B, compare to banding pattern in Fig 4B). Furthermore, the ΔPTM signal appeared lower on the western blot than the WT, as we have noted previously (compare to bands in Fig 4B). Taken together, these data suggest that both WT and ΔPTM protein VII are acetylated during infection, suggesting that mutation of known sites to prevent modification may result in acetylation at other sites.

**Figure 5:**
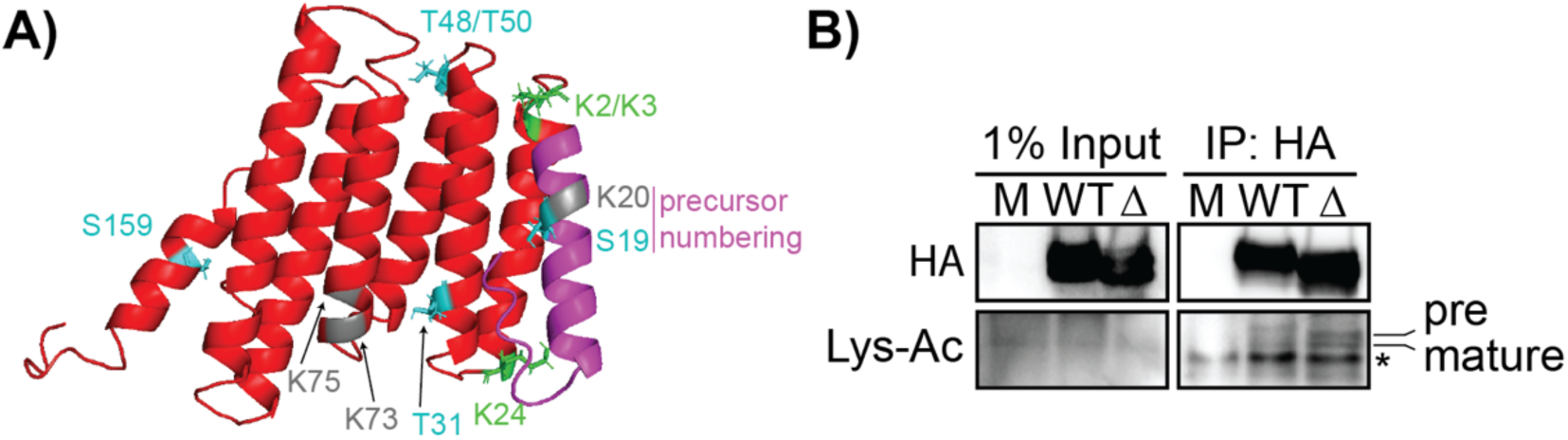
VIIΔPTM is acetylated at alternative residues in infected cells. A) I-TASSER structural prediction of protein VII. Protein VII is shown in red, pre-peptide in magenta, acetylation sites in green, and phosphorylation sites in cyan. Potential lysine residues that could be acetylated in protein VII are in dark gray. All modified or potentially modified residues are numbered with location of residue within mature protein, except S19 and K20 which are numbered within the precursor fragment. B) Western blot of protein VII immunoprecipitation (IP) at 24 h p.i. for mock, WT, and ΔPTM viruses as indicated. Blots were stained for HA (protein VII) and acetylated lysine. Bands for pre-VII and mature VII are labeled, * denotes the light chain of the IgG1 antibody used for IP.

### Protein VII’s interaction with E1A is chromatin dependent, and PTM sites are dispensable for this interaction

Next, we hypothesized that the PTMs may impact protein VII’s interactions with other proteins that may affect infection progression. We previously determined that protein VII directly interacts with host chromatin factor HMGB1 and that the ΔPTM mutant has a weaker interaction with HMGB1 (35). Protein VII was reported to interact with the early viral protein E1A and is hypothesized to recruit newly synthesized E1A to the viral genome during early infection (25). Because the K2AK3A virus appeared to have enhanced viral genome entry and faster early gene expression during early stages of infection (Fig 2 and 3), we hypothesized that PTMs may be important for protein VII’s interaction with newly expressed E1A on the viral genome, thus influencing early viral gene transcription. To test whether the two proteins directly interact, we performed a bacterial two-hybrid (B2H) analysis. Surprisingly, neither WT nor ΔPTM showed a positive interaction with E1A in our B2H, suggesting that they do not interact directly (Fig 6A).

**Figure 6:**
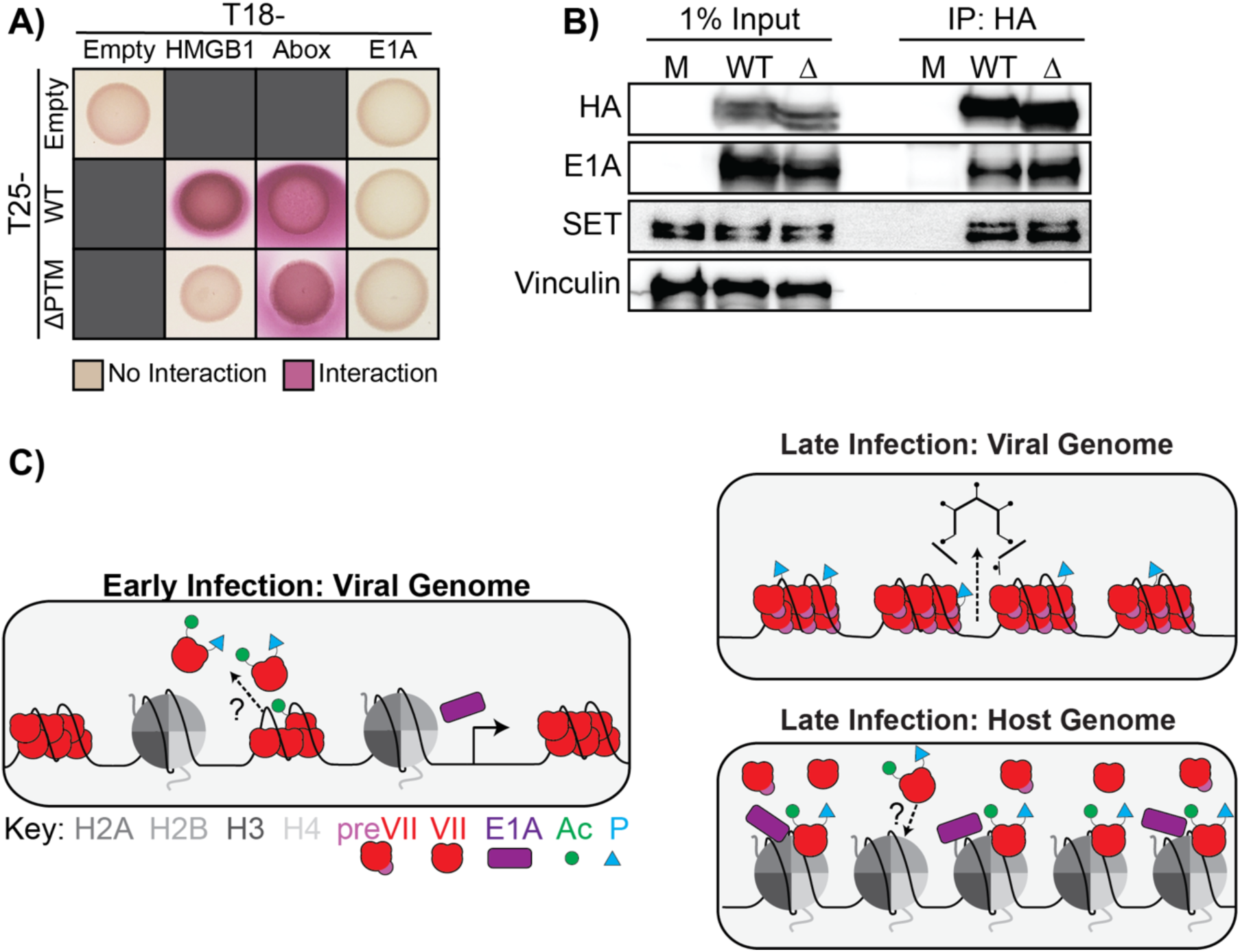
Protein VII-E1A interaction is chromatin dependent, and modified residues on protein VII are dispensable for this interaction. A) Bacterial two hybrid results of the interaction between E1A and WT and ΔPTM protein VII. Controls include empty vector (negative control), HMGB1 (positive control), or the A-box of HMGB1 (positive control). A positive interaction results in a pink color while a negative interaction remains beige. Representative image of 4 biological replicates. B) Representative western blot results from immunoprecipitation of HA from A549 cells infected with WT or ΔPTM viruses at 24 h p.i. Blots were stained for HA (protein VII), E1A, SET (positive control), and vinculin (negative control). n = 3 biological replicates. Note: HA IP is the same as in Fig 5B. C) Model of the role of PTMs on protein VII during early and late adenovirus infection. Within the virion, the viral genome is bound by unmodified or minimally modified protein VII. During early infection, the viral genome and protein VII are delivered to the nucleus. Protein VII is removed from the viral genome and replaced by host nucleosomes through an unknown mechanism. When PTMs are deposited on protein VII is unknown, although their addition may be important for accumulation of the viral genome in the nucleus and removal of protein VII from the viral genome. E1A interacts with protein VII in a chromatin-dependent manner and is recruited to the viral genome where it utilizes host transcription factors to initiate transcription of other early viral genes. During late infection, newly synthesized protein VII either localizes to host chromatin or is packaged with the viral genome within viral progeny. Modified protein VII is hypothesized to interact with host chromatin, whereas unmodified or minimally phosphorylated protein VII is packaged with the viral genome in progeny virions. Furthermore, while the difference in localization of preVII and mature protein VII is unknown, preVII is packaged in the virion as it is cleaved by the viral protease within the virion.

Because the B2H setting does not contain chromatin or other cellular factors, we sought to recapitulate the reported interaction in cells in the context of adenovirus infection. We infected A549 cells with the WT and ΔPTM viruses and assessed the interaction between protein VII and E1A by IP at 24 h p.i. In contrast to our B2H results, we observed that E1A robustly co-immunoprecipitated with WT protein VII, indicating the two proteins interact in cells during infection (Fig 6B). Furthermore, because protein VII ΔPTM also co-immunoprecipitated with E1A, this suggests that the mutated sites are not critical for this interaction. These results indicate that while there is likely no direct interaction between protein VII and E1A, because both proteins are found on chromatin throughout infection their interaction is likely mediated by chromatin. Taken together, these findings suggest that it may be through interacting with chromatin, viral or host, that protein VII and E1A come into contact.

## Discussion

In this study, we sought to determine how PTMs on the histone-like protein VII affect its function and in turn adenovirus infection. To our knowledge, this is the first attempt to characterize the function of these PTMs on protein VII during infection. We found that when acetylation of K2 and K3 of protein VII was prevented, adenovirus nuclear entry and early gene expression were both enhanced (Fig 2 and 3). Changes in early gene expression did not impact the later stages of infection as genome replication, late gene expression, and progeny production were comparable to wild type virus infection (Fig 4). Furthermore, mutating protein VII to mimic or prevent modification during infection had no effect on protein VII localization late in infection (Fig 4D). Interestingly, we also found that protein VIIΔPTM is likely acetylated at alternative residues to compensate for the introduced mutations (Fig 5).

What is the significance of acetylation on protein VII? In the literature, there are conflicting reports on whether protein VII is an activator or repressor of the initial burst of viral transcription (17, 18, 22, 25, 29–32). Reported results may be difficult to interpret due to different methods used with the complicating layer of regulation dictated by protein VII PTMs. Our results indicate that acetylation at the K2/K3 locus on protein VII is important for viral genome entry to the nucleus as loss of this site led to enhanced nuclear entry and earlier viral gene expression. Because this acetylation was identified upon ectopic expression, that is with no other viral proteins present, it is likely that a host protein is depositing the acetyl mark. Taken together, these observations suggest that a host acetyltransferase may be acting in a defense mechanism to delay viral entry. Nevertheless, the virus is able to overcome this barrier as the loss of these sites results in later viral infection dynamics that are indistinguishable from wild type. It is possible that enhanced viral genome entry is not due to the lack of acetylation but rather the change in charge. Our finding that protein VIIΔPTM is still acetylated despite the mutations suggests that either acetylation is critical for protein VII function, or the potential host defenses are ineffective. Future work to identify the enzyme or enzymes responsible will shed light on whether innate immune signals are responsible.

We found that protein VII’s interaction with the early viral protein, E1A, is mediated by chromatin and that PTM sites on protein VII are dispensable for this interaction (Fig 6). It is well-established that the first gene expressed upon entry of the viral genome into the nucleus is E1A. Newly synthesized E1A promotes further viral transcription on active viral genomes while also binding to the host genome to influence multiple pathways for viral benefit (26, 43, 44).

Evidence for the interaction of protein VII with E1A led to a model in which the newly translated E1A localizes to the viral genome by binding directly to protein VII, which is still present on the incoming viral genomes (25). Our results in this study suggest that E1A binds to protein VII only in a chromatin-mediated manner (i.e. in eukaryotic cells but not bacteria, see Fig 6), which would suggest that the order of events is for (1) histones to be deposited on the viral genome together with protein VII, and then (2) E1A to bind and activate expression of other viral genes. While we cannot rule out other models, we propose that protein VII promotes the remodeling of the incoming viral genome to encourage deposition of host histones, together with TAF-1β/SET (18, 22), which in turn allow for the expression of E1A, that then binds to these genomes and promotes further viral gene expression (Fig 6C). On the host genome, E1A has been well-established to alter histone acetylation and promote an S-like phase. Protein VII also localizes to the host genome, although whether protein VII and E1A localize to the same host genomic loci is unknown. Future work with genomic profiling methods will elucidate the overlap of these two viral proteins on host chromatin and the direct outcome on host transcription.

The K2/K3 site of modification in the wild-type virus is an AKKRS motif, which is reminiscent of the canonical histone ARSK motif. Histone mimics have been observed on Influenza A Virus H3N2 NS1 protein and SARS-CoV-2 ORF8, where viral proteins mimic the histone H3-tail to subvert immune responses (45–47). In the case of protein VII, it is possible that this AKKRS motif is a viral mimic of the histone motif (48, 49). Protein VII is highly conserved amongst human adenoviruses, and the histone mimic motif is also conserved across HAdV species as well as within other vertebrate adenoviruses, including an ARKRS motif in mouse MAdV-1. Despite the conservation of this motif, protein VII in general varies widely across different adenovirus species in sequence and length. Surprisingly, protein VII from reptilian and avian adenoviruses are only 128 and 78 amino acids in length, respectively (50, 51). While the origins of protein VII are not known, given its generally poor conservation across non mammalian species, protein VII is likely to be rapidly evolving such that mimicking histones may be a recent adaptation. Future work into the evolution of protein VII will shed more light on the importance of the conservation and function of potential PTMs at these sites.

We found that increased early gene expression had no significant impact on later stages of infection or the production of infectious progeny (Fig 4). We also found that the ΔPTM protein VII was still acetylated during infection, suggesting that other sites on protein VII may be modified to compensate for the mutations we introduced (Fig 5). We observed the distinctive double band of protein VII in the WT and ΔPTM samples, suggested that the both the preVII and mature VII proteins are acetylated in ΔPTM. Thus, either K73 or K75 in mature protein VII is likely acetylated since there are no other lysine residues within mature protein VIIΔPTM. In contrast, there are twelve threonine and six serine residues throughout protein VII (e.g. S5, T28, T48, S130, S161, T169). Thus, it is possible that mutation of the three phosphorylated sites in ΔPTM may not produce an observable defect in the virus due to compensation by phosphorylation of another site. Another possibility is that phosphorylation of various serine and threonine residues within protein VII may be non-specific and dynamic. Future studies employing kinase screens may be effective at identifying potential enzymes or pathways responsible for protein VII phosphorylation and how phosphorylation of protein VII impacts infection.

During infection, protein VII is first expressed as a precursor protein, preVII, which is cleaved by a viral protease to produce the mature protein (12, 13), the latter of which has been the focus of this study. Because the prior ectopic expression analyses were done with the mature protein only, it is possible that the presence of the precursor peptide may mask the effects of the PTMs. Indeed, the precursor fragment contains both a nucleolar localization and reported PTMs (52, 53), which may impact its function. While the role of the precursor is not well understood beyond virion packaging, the ratio of precursor to mature protein heavily favors the precursor based on western blotting during late infection. Furthermore, the exact amount of mature protein VII in the nucleus that is not packaged within progeny virions is unknown. Taken together, these observations indicate that further study is needed to elucidate the functional differences between the mature and precursor proteins.

## Acknowledgements

We would like to thank the Avgousti lab and M. Weitzman for helpful comments and critical insight. We also thank R. Strong for advice with I-TASSER protein structure prediction. This study was supported by NIH funding to E.A.A. (T32AI083203), J.G.S. (R01AI104920), M.S.G. (R00GM134153), and D.C.A. (R35GM133441), the University of Washington Helen Riaboff Whiteley Fellowship to E.A.A. and Magnuson Scholarship to L.E.K.M., and startup funds from the University of Washington (M.S.G., https://microbiology.washington.edu/) and Fred Hutchinson Cancer Center (D.C.A., https://www.fredhutch.org/en.html).

## Methods and Materials

### Cell lines, viruses, and infections

A549 cells were purchased from ATCC and cultured in Kaighn’s modification of Ham’s F-12 medium (F-12K) containing 100 U/ml of penicillin and 100 mg/ml of streptomycin and supplemented with 10% fetal bovine serum (FBS). HEK293T cells were purchased from ATCC and HEK293Q cells were purchased from Qiagen. Both were cultured in Dulbecco’s Modified Eagle Medium (DMEM) containing 100 U/mL of penicillin and 100 mg/ml of streptomycin and supplemented with 10% fetal bovine serum (FBS). HeLa cells were purchased from ATCC and cultured in Eagle’s Minimum Essential Medium (EMEM) containing 100 U/mL of penicillin and 100 mg/ml of streptomycin and supplemented with 10% fetal bovine serum (FBS).

Ad5 VIIΔPTM-HA, VIIK2AK3A-HA, and VIIK3Q-HA were all generated by recombineering using a bacterial artificial chromosome containing the genome of a replication-competent, E3-deleted HAdV-5-based vector containing an HA-tagged protein VII (35, 38). Successful recombineering was verified through restriction digest and Sanger sequencing. To produce virus, viral genomes were linearized by *Pac* I endonuclease digestion and transfected into 293β5 cells (54). The resulting virus was amplified by repeated passaging in 293β5 cells (54), purified with two rounds of ultracentrifugation in a CsCl gradient, snap frozen in liquid nitrogen, and then stored in a 20% glycerol buffer at −80°C (55). Viral titers were determined by flow cytometry and plaque assay.

Infections for figures 1-2 were performed at a multiplicity of infection of 10 or 25 plaque forming units per cell. Infections were synchronized by adding inoculum to cells on ice and incubating for 1 h while rocking every 20 min. Cells were moved to 37°C for an additional 2 h with rocking every 20 min before the inoculum was removed and replaced with fresh media. Infections for Figures 3-5 were carried out likewise but without synchronization on ice.

### Plaque assays

Cell pellets from time course infections of A549 cells were collected at 24, 36, and 48 h p.i. Cell pellets were resuspended in 100 µL PBS, freeze-thawed 4x, and pelleted in a table-top centrifuge at max speed, and virus containing supernatant was collected for plaque assays. Plaque assays were performed on HEK293Q cells. Cells were seeded in 6-well plates, infected the next day with serial dilutions of virus samples, and overlayed with a 4% SeaPlaque agarose. Plates were incubated at 37 °C until plaques developed, approximately 6-7 days. The agarose overlay was dissolved by incubating with 10% trichloroacetic acid in phosphate buffered saline at RT for 30 min and then stained with crystal violet. Plaques were counted by eye, and the count was used to determine the concentration of infectious units.

### Antibodies

Commercially available antibodies were purchased through Abcam (HMGB1 [18526], H3 [ab1791], HA-tag [ab9110], SET [ab181990], Adenovirus Late Proteins [ab6982]), Sigma-Aldrich (Vinculin [V9131]), and BD Biosciences (E1A [554155]). Protein VII antibodies were a generous gift from the Gerace and Wodrich labs. DBP antibodies were a generous gift from the Levine lab. Secondary antibodies used for immunoblotting were obtained from Jackson ImmunoResearch (115-035-003 and 111-035-045). Secondary antibodies for immunofluorescence microscopy were obtained from Thermo Fisher Scientific (A-11011, A-11001). DAPI stain was obtained from Fisher Scientific (50-874-10001).

### Immunofluorescence microscopy

Immunofluorescence microscopy was performed as previously described (34, 35). Briefly, cells were seeded on poly-L-lysine coated coverslips in a 24-well plate. Cells were infected with viruses, and coverslips were fixed at 0, 2, 4, 6 and 8 hh p.i. with 4% paraformaldehyde. After fixation, cells were permeabilized with 0.5% Triton-X, washed three times with phosphate buffered saline (PBS), and blocked with 3% BSA. Cells were incubated with primary antibodies for 1 h at RT, washed three times in PBS, incubated with secondary antibody and DAPI for 1 h in the dark, and then washed three times in PBS. Coverslips were then mounted on slides with ProLong Gold Antifade Mountant (Thermo Fisher Scientific) and allowed to dry overnight. High resolution confocal microscopy was performed with a Leica Stellaris Confocal Microscope using a 63x oil objective.

### RT-qPCR and qPCR

For qPCR, gDNA was extracted from cells with Qiagen QIAamp DNA kit. Extracted gDNA was normalized to 50 ng/µL and then used for qPCR with primers targeting viral DBP and cellular tubulin for normalization. qPCR results in Fig 2C were determined by performing a standard curve with viral DBP primers against a BAC containing the HAdV-5 VII-HA viral genome. The standard curve was used to determine ng of input DNA for each virus, which was converted to the number of total genomes, and then divided by the number of cells present to quantify the number of genomes per cell. qPCR results in Fig 4 are depicted as fold-change over an input control at 4 h p.i.

For RT-qPCR, RNA was extracted from cells with Trizol. Extracted RNA was converted to cDNA with Iscript Reverse Transcription Supermix (BioRad). cDNA at a concentration of 50 ng/µl was used for qPCR with iTaq Universal SYBR Green Supermix (BioRad) with primers spanning an exon-intron boundary of E1A and RPLP0 for normalization (see primers table below). RT-qPCR results are depicted as fold-change over internal control gene RPLP0. For transcriptional and replication analysis, qPCR was performed with the BioRad CFX384 Real-Time System.

### Western blotting

Samples were resuspended in 1X Laemmli sample buffer with 5% β-mercaptoethanol (200µL per 10^6^ cells), separated on 12% or 15% polyacrylamide gels, and then transferred to nitrocellulose. Membranes were blocked in 5% milk in TBST buffer or 5% BSA in TBST buffer for 30 min, and then probed with primary antibodies at 4°C overnight. Blots were then washed with TBST buffer for 30 min, probed with HRP-conjugated secondary antibodies for 1 h at RT, washed with TBST buffer for 30 min, and developed with Clarity Western ECL Substrate, and imaged with a Biorad ChemiDoc MP Imaging System.

### Protein prediction

Amino acid sequence of wild-type protein VII (NCBI accession ID AAW65510) was run on the protein prediction software I-TASSER (41, 42). The top predicted model was selected, and the 3D model was formatted in PyMOL.

### Bacterial 2-hybrid

Plasmids (pUT18, pUT18C, pKT25, and pKNT25) containing fusion constructs of HMGB1, A-box, protein VII, and E1A were co-transformed into the B2H assay strain BTH101. Four replicates from each transformation were picked, grown in M63 minimal media for 48 h, and spotted on MacConkey agar plates supplemented with carbenicillin, kanamycin, 1% maltose, and 1mM IPTG. Plates were incubated at 30°C and imaged after 72 hrs. Interaction between protein VII and E1A (e.g., T25-VII with T18-E1A, T25-VII with E1A-T18, VII-T25 with T18-E1A, and VII-T25 with E1A-T18).

### Immunoprecipitation

A549 cells were infected with Ad5 VII-HA or Ad5 VIIΔPTM-HA at an MOI of 10. Cell pellets were collected at 24h p.i. and stored at −80°C until ready to proceed to IP. Cells were thawed on ice, resuspended in 1mL of lysis buffer (20mM Hepes pH 7.4, 110mM KOAc, 2mM MgCl_2_, 0.1% Tween-20. 0.5% Triton-X 100, 200mM NaCl, protease/phosphatase inhibitors (Thermo Fisher A32961, added fresh before use), and incubated on ice for 10 min mixing intermittently. The lysate was then treated with 5µL of benzonase (Fisher Scientific 71-205-3) for one h at 4°C with rocking. Lysates were pelleted at 4°C at max speed for 15 min. The supernatant was transferred to new tubes and then protein concentration from each sample was measured by Bradford assay. Samples were normalized to 1 ml of 2mg/mL protein, and 100µL was removed as 10% input sample. 60µL of HA-conjugated beads was washed twice with lysis buffer and then ∼20µL of beads was added to remaining 900µL of protein samples. Samples were incubated with rocking at 37°C. Beads were washed two times with 1 mL of lysis buffer, and then eluted with 100µL of 1X sample buffer (10% DTT) at 95°C for 20 min. Eluted samples were then separated from beads with a magnetic stand. 30µL of IP sample and 10µL of input sample (1%) were run on an SDS-PAGE gel for western blotting.

### Statistical analyses

All statistical analyses were performed using GraphPad Prism v10. Statistical tests and n values are described in the figure legends. Statistical significance was defined as p<0.05 in all experiments. Specifically, we used a one-way ANOVA with Dunnett’s multiple comparisons test or a two-way ANOVA with a Tukey’s multiple comparisons test where described. Only p-values less than cutoff are reported in figures.

### Primers

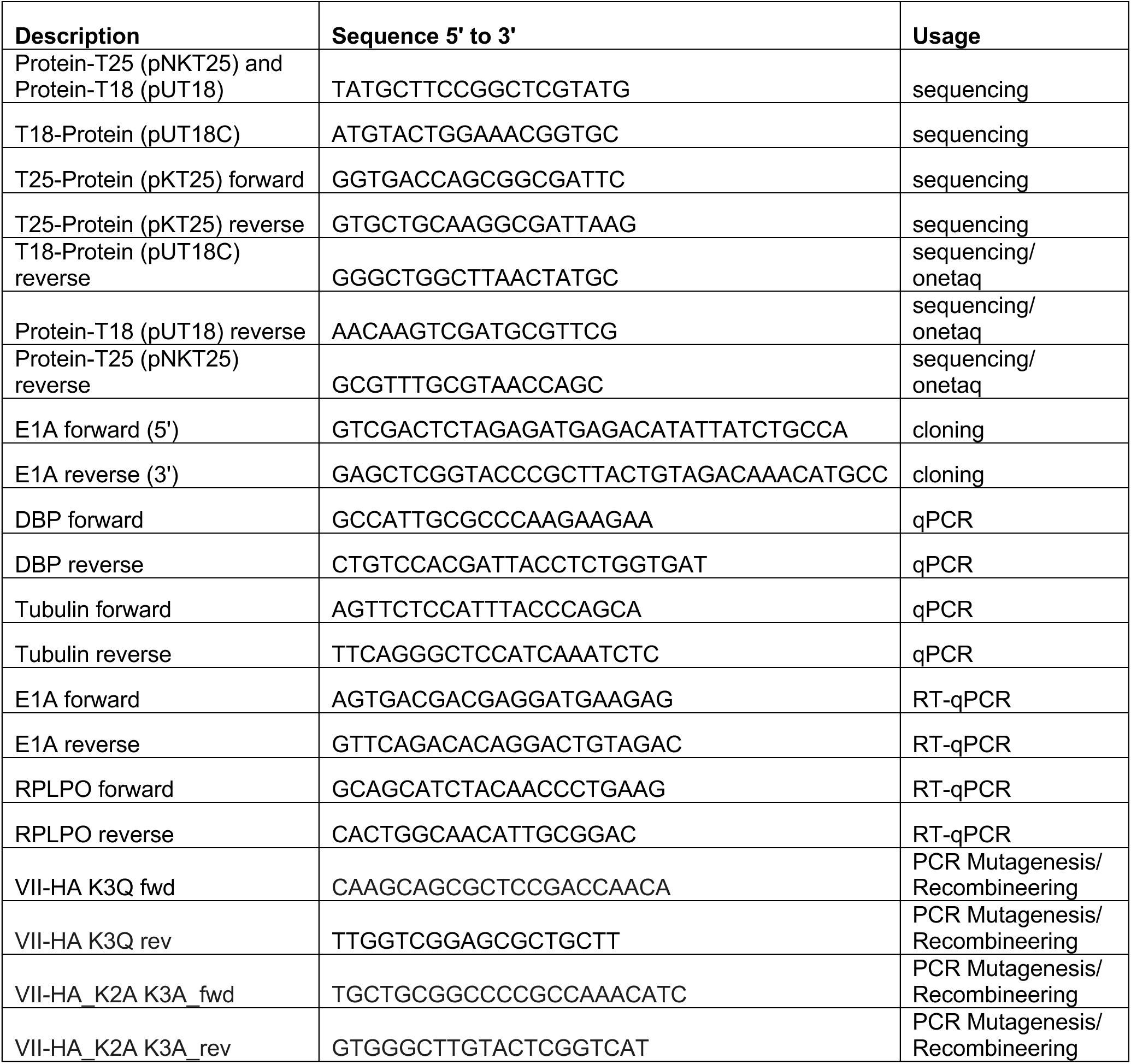

